## Supplementary material for "Soil microbiota and herbivory drive the assembly of plant-associated microbial communities through different mechanisms"

### Contents

**Supplementary figures** . . . . . . . . . . . . . . . . . . . . . . . . . . . . . . . . . . . . . **1**

**Supplementary tables** . . . . . . . . . . . . . . . . . . . . . . . . . . . . . . . . . . . . . **6**

### Supplementary figures

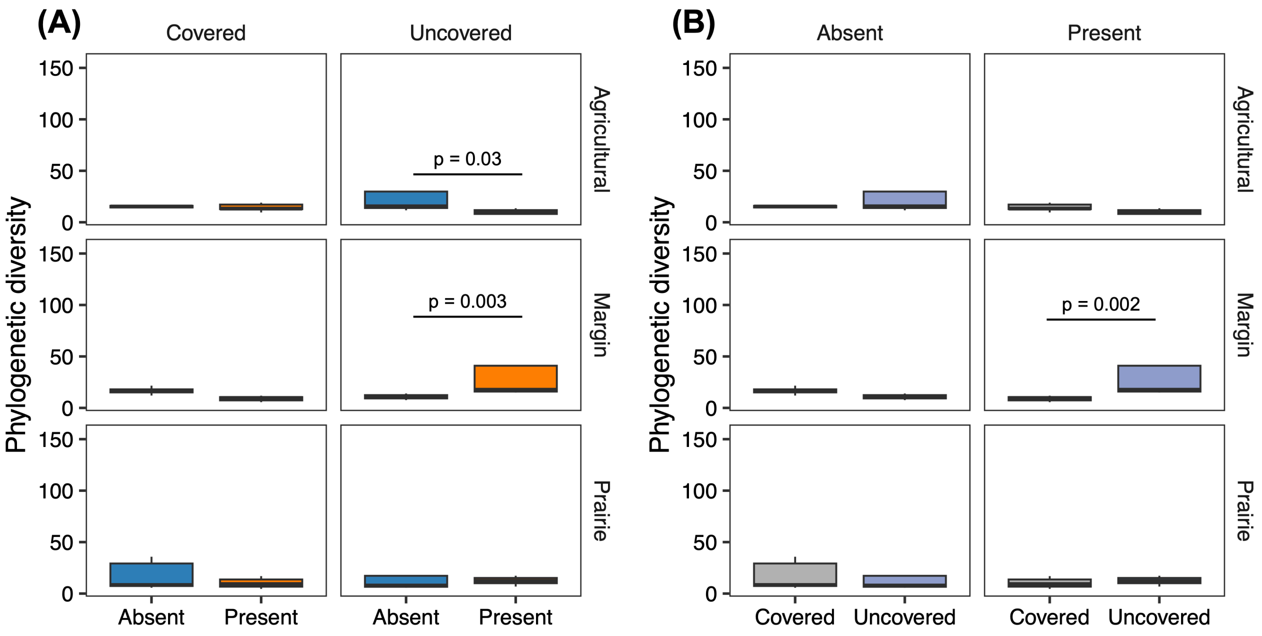


**Figure S1**. Phylogenetic diversity in samples in the rhizosphere compartment. **(A)** Comparison between samples exposed to herbivory (present) and control (absent) for each combination of soil inoculum (agricultural, margins, prairie) and coverage (covered, uncovered). **(B)** Comparison between samples from microcosms with covered and uncovered surface, for each combination of soil inoculum (agricultural, margins, prairie) and herbivory (present, absent). Post-hoc p-values are FDR-corrected.

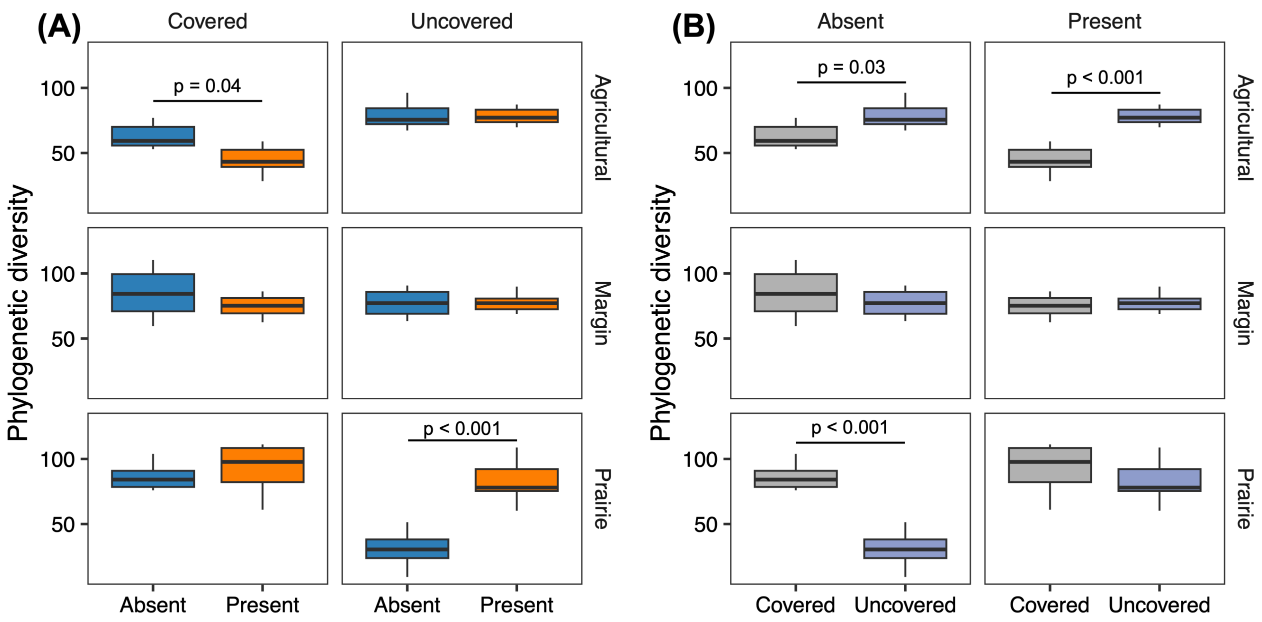


**Figure S2**. Phylogenetic diversity for samples in the root compartment. **(A)** Comparison between samples exposed to herbivory (present) and control (absent) for each combination of soil inoculum (agricultural, margins, prairie) and coverage (covered, uncovered). **(B)** Comparison between samples from microcosms with covered and uncovered surface, for each combination of soil inoculum (agricultural, margins, prairie) and herbivory (present, absent). Post-hoc p-values are FDR-corrected.

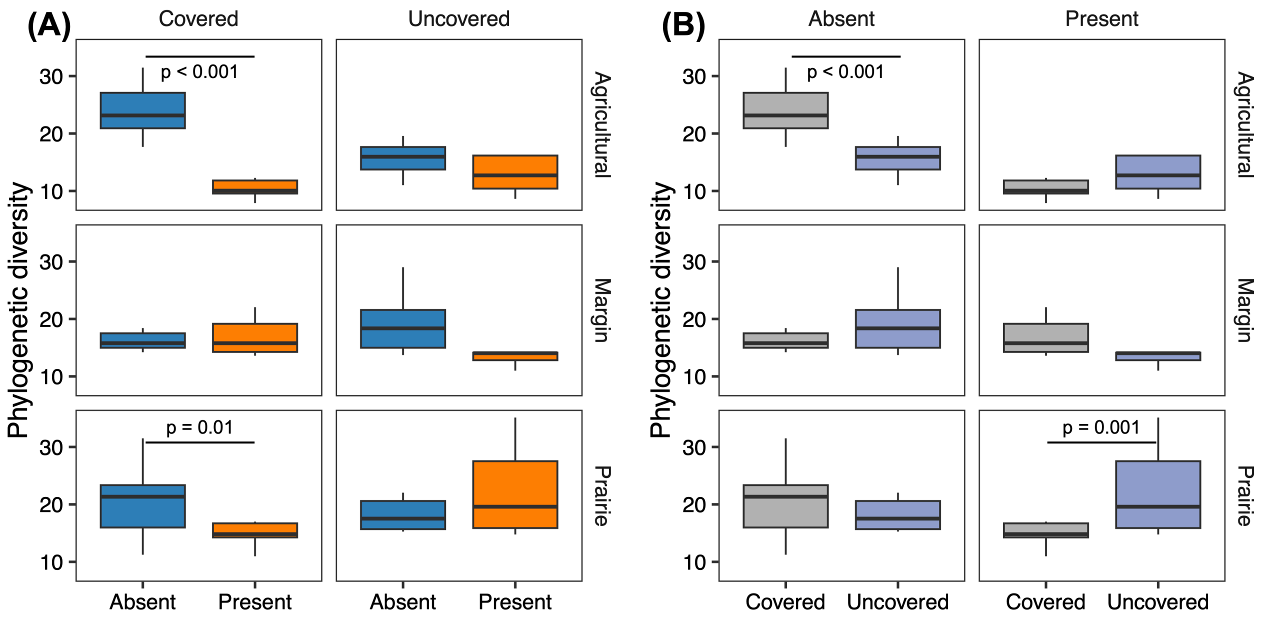


**Figure S3**. Phylogenetic diversity for samples in the leaf compartment. (A) Comparison between samples exposed to herbivory (present) and control (absent) for each combination of soil inoculum (agricultural, margins, prairie) and coverage (covered, uncovered). (B) Comparison between samples from microcosms with covered and uncovered surface, for each combination of soil inoculum (agricultural, margins, prairie) and herbivory (present, absent). Post-hoc p-values are FDR-corrected.

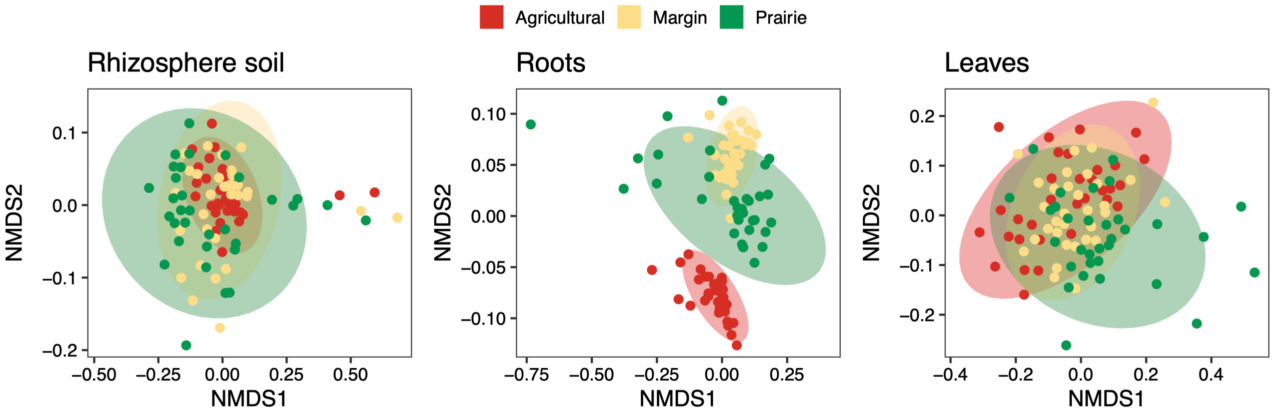


**Figure S4**. NMDS plots of bacterial community Unifrac distance matrix for each compartment. Points and 95% CI ellipses are coloured by soil inoculum.

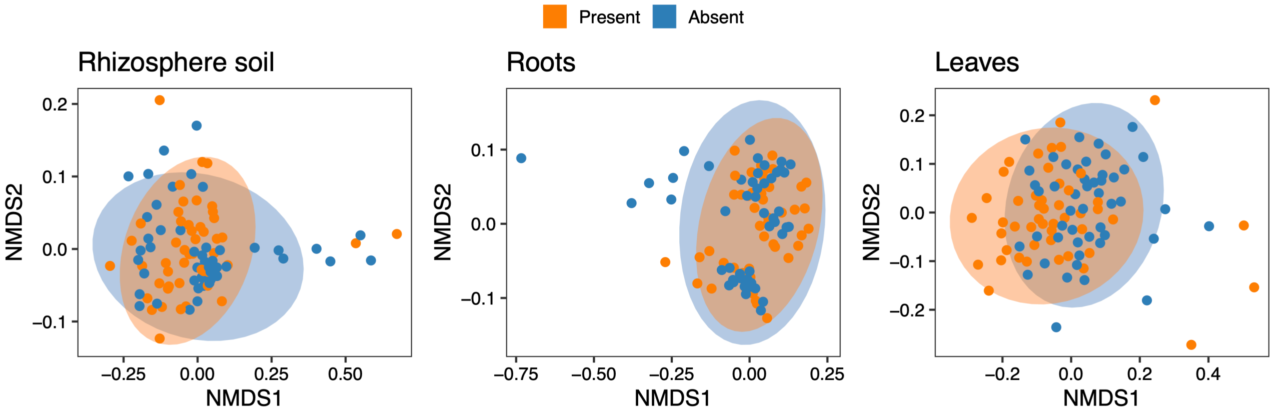


**Figure S5**. NMDS plots of bacterial community Unifrac distance matrix for each compartment. Points and 95% CI ellipses are coloured by presence or absence of herbivore.

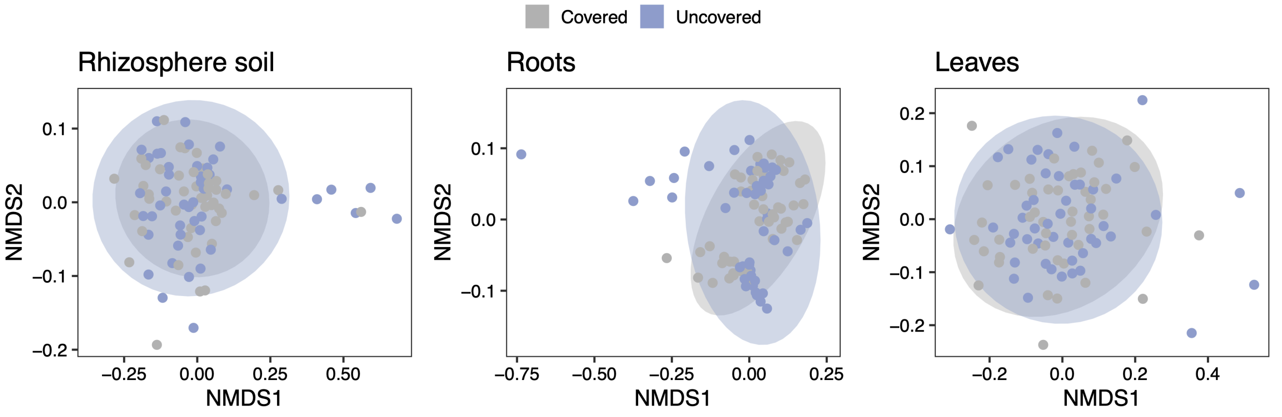


**Figure S6**. NMDS plots of bacterial community Unifrac distance matrix for each compartment. Points and 95% CI ellipses are coloured by presence or absence of soil cover.

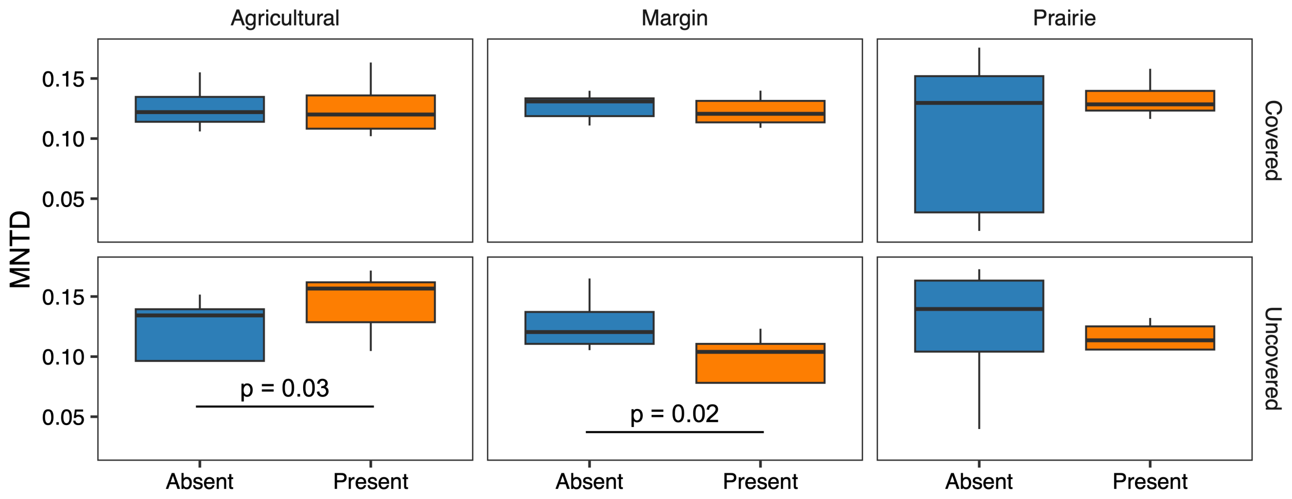


**Figure S7**. Mean Nearest Taxon Distance (MNTD) for samples in the rhizosphere compartment. Comparison between samples from microcosms with herbivores or control group, for each combination of soil inoculum (agricultural, margins, prairie) and cover treatment (covered, uncovered). Post-hoc p-values are FDR-corrected.

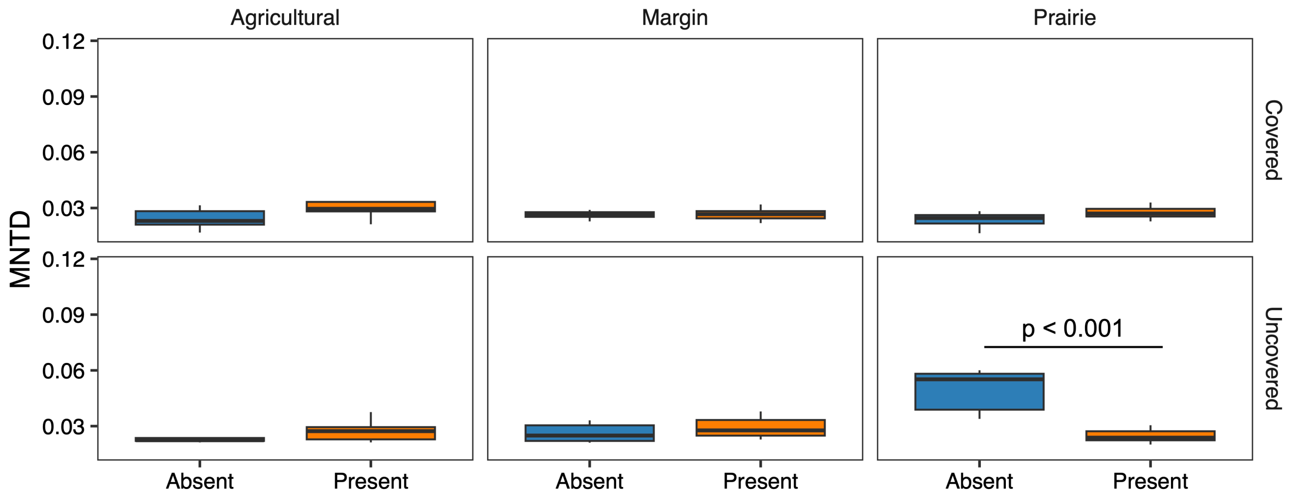


**Figure S8**. Mean Nearest Taxon Distance (MNTD) for samples in the root compartment. Comparison between samples from microcosms with herbivores or control group, for each combination of soil inoculum (agricultural, margins, prairie) and cover treatment (covered, uncovered). Post-hoc p-values are FDR-corrected.

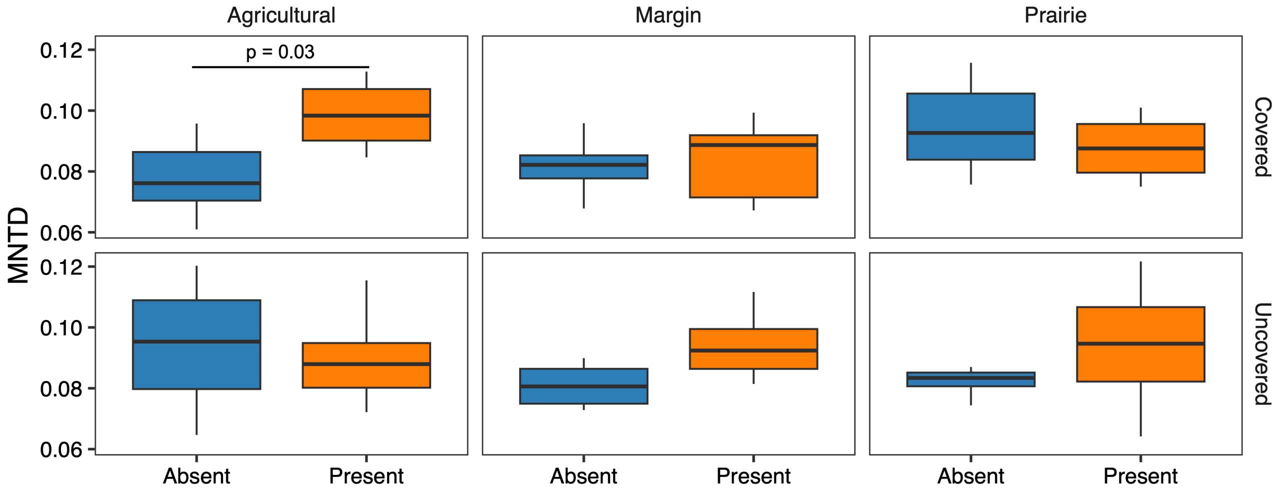


**Figure S9**. Mean Nearest Taxon Distance (MNTD) for samples in the leaf compartment. Comparison between samples from microcosms with herbivores or control group, for each combination of soil inoculum (agricultural, margins, prairie) and cover treatment (covered, uncovered). Post-hoc p-values are FDR-corrected.

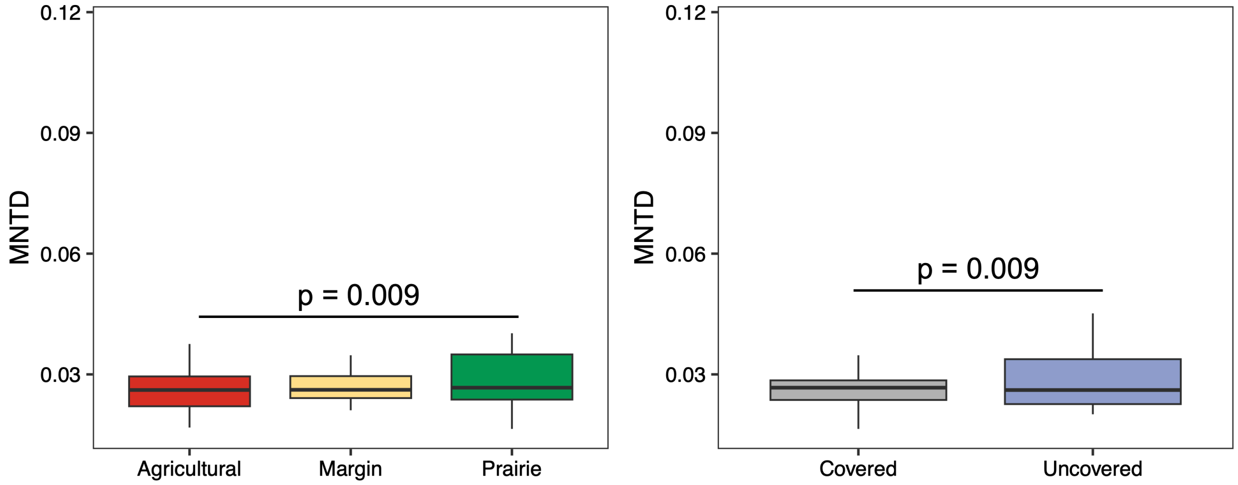


**Figure S10**. Mean Nearest Taxon Distance (MNTD) for samples in the root compartment. **(A)** Comparison between samples from microcosms with different soil inocula (agricultural, margins, prairie) and **(B)** cover treatment (covered, uncovered). Post-hoc p-values are FDR-corrected.

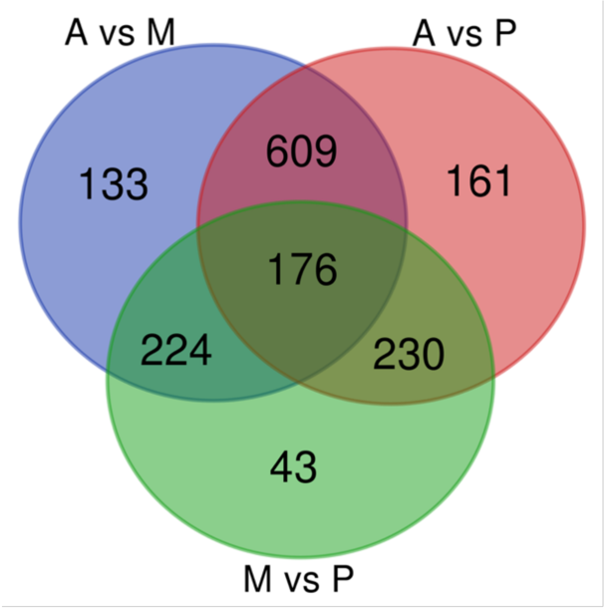


**Figure S11**. Venn diagram comparing ASVs which abundance was influenced by the soil inoculum across all pairwise comparisons: agricultural soil vs field margins (A vs M), agricultural soil vs prairie soil (A vs P), field margins vs prairie soil (M vs P).

### Supplementary tables

**Table S1**. Results from a linear mixed-effects model for each compartment (rhizosphere soil, roots, leaves, herbivore) testing the effect of soil inoculum (S; agricultural, margins, prairie), herbivory (H; present, absent), coverage (T; present, absent), and all their interactions on the phylogenetic diversity index. The variance explained ($R^2$) is reported for each of the single factor.

| | | **Rhizosphere** | | | **Roots** | | | **Leaves** | | | **Herbivore** | | |
|---|---|---|---|---|---|---|---|---|---|---|---|---|---|
| **Factor** | df | $\chi^2$ | p | $R^2$ | $\chi^2$ | p | $R^2$ | $\chi^2$ | p | $R^2$ | $\chi^2$ | p | $R^2$ |
| **Soil (S)** | 2 | 0.51 | 0.77 | 0.005 | 8.4 | 0.01 | 0.038 | 9.17 | 0.01 | 0.056 | 7.31 | 0.02 | 0.149 |
| **Herbivory (H)** | 1 | 0.48 | 0.48 | 0.004 | 4.39 | 0.03 | 0.024 | 14.47 | 0.001 | 0.092 | | | |
| **Coverage (C)** | 1 | 2.13 | 0.14 | 0.019 | 1.25 | 0.26 | 0.005 | 0.03 | 0.84 | 0.0002 | 0.07 | 0.78 | 0.001 |
| **S × H** | 2 | 6.81 | 0.03 | | 32.49 | 0.001 | | 13.98 | 0.001 | | | | |
| **S × C** | 2 | 2.5 | 0.28 | | 56.26 | 0.001 | | 6.4 | 0.04 | | 0.89 | 0.63 | |
| **H × C** | 1 | 1.13 | 0.28 | | 17.34 | 0.001 | | 8.15 | 0.004 | | | | |
| **S × H × C** | 2 | 8.46 | 0.01 | | 6.18 | 0.04 | | 16.58 | 0.001 | | | | |

**Table S2**. PERMANOVA post-hoc pairwise comparison between covered and control groups for each herbivory x soil inoculum combination, for both rhizosphere soil and roots bacterial microbiota. Values represent FDR-corrected p-values.

| | | Rhizosphere soil | Roots | Leaves |
|---|---|---|---|---|
| **Herbivory** | *Agricultural* | 0.026 | 0.001 | 0.012 |
| | *Margin* | 0.004 | 0.037 | 0.022 |
| | *Prairie* | 0.166 | 0.001 | 0.005 |
| **Control** | *Agricultural* | 0.005 | 0.001 | 0.005 |
| | *Margin* | 0.002 | 0.047 | 0.066 |
| | *Prairie* | 0.271 | 0.039 | 0.203 |

**Table S3**. PERMANOVA post-hoc pairwise comparison between herbivory and control groups for each coverage x soil inoculum combination, for both rhizosphere soil and roots bacterial microbiota. Values represent FDR-corrected p-values.

| | | Rhizosphere soil | Roots | Leaves |
|---|---|---|---|---|
| **Covered** | *Agricultural* | 0.004 | 0.008 | 0.003 |
| | *Margin* | 0.002 | 0.009 | 0.012 |
| | *Prairie* | 0.045 | 0.029 | 0.003 |
| **Control** | *Agricultural* | 0.004 | 0.019 | 0.006 |
| | *Margin* | 0.002 | 0.804 | 0.127 |
| | *Prairie* | 0.046 | 0.001 | 0.1 |

**Table S4**. Results from a linear mixed-effects model for each compartment (rhizosphere soil, roots, leaves, herbivore) testing the effect of soil inoculum (S; agricultural, margins, prairie), herbivory (H; present, absent), coverage (T; present, absent), and all their interactions on the MNTD index.

| | | Rhizosphere | | Roots | | Leaves | | Herbivore | |
|---|---|---|---|---|---|---|---|---|---|
| **Factors** | **df** | $\chi^2$ | **p** | $\chi^2$ | **p** | $\chi^2$ | **p** | $\chi^2$ | **p** |
| **Soil (S)** | 2 | 1.15 | 0.56 | 10.85 | 0.004 | 2.8 | 0.24 | 1.98 | 0.37 |
| **Herbivory (H)** | 1 | 0.21 | 0.64 | 1.57 | 0.21 | 3.81 | 0.05 | | |
| **Coverage (C)** | 1 | 0.42 | 0.51 | 7.07 | 0.007 | 0.19 | 0.66 | 0.16 | 0.68 |
| **S × H** | 2 | 6.01 | 0.04 | 17.81 | 0.001 | 0.9 | 0.63 | | |
| **S × C** | 2 | 2.26 | 0.32 | 15.57 | 0.001 | 1.08 | 0.58 | 1.54 | 0.46 |
| **H × C** | 1 | 0.71 | 0.39 | 12.16 | 0.001 | 0.18 | 0.66 | | |
| **S × H × C** | 2 | 6.64 | 0.03 | 18.95 | 0.001 | 7.93 | 0.01 | | |

**Table S5**. Results from chi-squared tests comparing the number of ASVs unique to a compartment or shared between compartments in plants grown on pots with covered and uncovered soil surface. See Figure 3.

| **Intersection** | **covered** | **uncovered** | $\chi^2$ | **p** |
|---|---|---|---|---|
| *herbivore* | 83 | 101 | 1.76 | 0.185 |
| *herbivore-leaves* | 1 | 2 | 0.33 | 0.564 |
| *herbivore-leaves-rhizosphere* | 2 | 3 | 0.20 | 0.655 |
| *herbivore-leaves-roots* | 95 | 19 | 50.67 | 0.001 |
| *herbivore-leaves-roots-rhizosphere* | 250 | 333 | 11.82 | 0.001 |
| *herbivore-rhizosphere* | 2 | 9 | 4.45 | 0.035 |
| *herbivore-roots* | 220 | 110 | 36.67 | 0.001 |
| *herbivore-roots-rhizosphere* | 87 | 262 | 87.75 | 0.001 |
| *leaves* | 48 | 54 | 0.35 | 0.552 |
| *leaves-rhizosphere* | 7 | 23 | 8.53 | 0.003 |
| *leaves-roots* | 148 | 89 | 14.68 | 0.001 |
| *leaves-roots-rhizosphere* | 66 | 164 | 41.75 | 0.001 |
| *rhizosphere* | 64 | 460 | 299.26 | 0.001 |
| *roots* | 3914 | 2821 | 177.37 | 0.001 |
| *roots-rhizosphere* | 183 | 1013 | 576.01 | 0.001 |

**Table S6**. ASVs in the root compartment significantly influenced by soil inoculum as pairwise contrasts between agricultural soil and field margins (sheet 1, A vs M), agricultural soil and prairie soil (sheet 2, A vs P), field margins and prairie (sheet 3, M vs P). Includes also ASVs in the root compartment significantly influenced by soil cover. **Table is attached as xlsx file**.
